# Database-Augmented Transformer-Based Large Language Models Achieve High Accuracy in Mapping Gene-Phenotype Relationships

**DOI:** 10.1101/2025.01.28.635344

**Authors:** Nicolas Matthew Suhardi, Anastasia Oktarina, Mathias P.G. Bostrom, Xu Yang, Vincentius Jeremy Suhardi

## Abstract

Transformer-based large language models (LLMs) have demonstrated significant potential in the biological and medical fields due to their ability to effectively learn from large-scale, diverse datasets and perform a wide range of downstream tasks. However, LLMs are limited by issues such as information processing inaccuracies and data confabulation, which hinder their utility for literature searches and other tasks requiring accurate and comprehensive extraction of information from extensive scientific literature. In this study, we evaluated the performance of various LLMs in accurately retrieving peer-reviewed literature and mapping correlations between 102 genes and four phenotypes: bone formation, cartilage formation, fibrosis, and cell proliferation. Our analysis included standard transformer-based LLMs (ChatGPT4o and Gemini1.5 Pro), fine-tuned LLMs with dedicated custom databases containing peer-reviewed articles (SciSpace and ScholarAI), and fine-tuned LLMs without dedicated databases (PubMedGPT and ScholarGPT). Using human-curated gene-to-phenotype mappings as the ground truth, we found that fine-tuned LLMs with dedicated databases (SciSpace and ScholarAI) achieved high accuracy (>80%) in gene-to-phenotype mapping. Additionally, these models were able to provide relevant peer-reviewed publications supporting each gene-to-phenotype correlation. These findings underscore the importance of database augmentation and finetuning in enhancing the reliability and utility of LLMs for biomedical research applications.

## Introduction

Understanding the relationship between genes and their cellular phenotypes is crucial in the biomedical field, as it provides insights into how genetic perturbations impact cellular behavior^1^. This knowledge is essential for determining how gene function relates to cellular phenotypes. The process of mapping genetic perturbations to phenotypes typically involves either identifying the gene(s) associated with a specific observed phenotype or evaluating the phenotypic changes resulting from the perturbation of specific gene(s). Accurate correlations between genes and phenotypes are foundational for various applications, including cellular type classification in single-cell omics analysis^2^. However, the traditional approach of perturbing one gene at a time is labor-intensive and time-consuming. Coupled with the exponential growth in the number of research publications^3^, this has made the task of accurately mapping genes to phenotypes increasingly challenging.

Large language models (LLMs), more specifically transformer-based LLM, have recently gained significant attention in the fields of biology and medicine due to their ability to effectively learn from large-scale, diverse datasets and their versatility in performing a variety of downstream tasks and often outperform task-specific models trained from scratch^4,5^. LLMs have proven to be powerful tools for deciphering and recapitulating the nuances and complex relationships within various types of genomics data (e.g., DNA sequences), transcriptomics data (e.g., RNA sequences), and proteomics data (e.g., protein sequences and structures)^6^. Transformer-based LLMs utilize the self-attention mechanism^7^ to understand context and handle long-range relationships in text, enabling them to perform tasks such as text translation, summarization, and generation much more efficiently than non-transformer-based models like recurrent neural networks (RNNs) and long short-term memory (LSTM) models^8^. Furthermore, the multi-head attention mechanism, an integral feature of the transformer architecture, allows transformers to process information in parallel, significantly reducing processing time compared to RNNs and LSTMs.^7^

Despite their potential and growing adoption in biomedical research applications, multiple studies have highlighted the downsides of LLMs, such as issues with accuracy^9^ and generation of false information (hallucination)^10^. Furthermore, lack of transparency in the data used for training and the bias in the training data can perpetuate errors^11^. With LLM being increasingly used in biomedical research to identify gene-phenotype relations^2,12^, there is a need to characterize LLM ability to accurately identify gene-phenotype relation.

In this study, we evaluate the degree to which transformer-based LLMs, such as ChatGPT-4o and Gemini 1.5 Pro, can identify gene-phenotype relationships based on their embedded biological knowledge and access to PubMed and other open access full-length articles. Using proliferation, osteogenesis, fibrogenesis, cartilage formation as proxies for phenotypes and using human-based peer-reviewed literature searches as the ground truth, ChatGPT-4o and Gemini 1.5 Pro were able to identify gene-phenotype relation with moderate accuracy. Further fine-tuning with abstracts and full-length articles significantly improves the accuracy of LLMs in identifying gene-phenotype relationships. Collectively, these findings suggest that fine-tuned LLMs with access to abstracts and full-text articles can serve as a reliable and accurate resource for mapping gene-phenotype relationships.

## Methods

### Transformer-based large language model

Publicly available transformer-based large language model ChatGPT-4o (“o” for “omni”) and Google Gemini 1.5 Pro were used through their user interface. As of January 2025, when prompted on the number of tokens and parameters, GPT-4, the foundational model underlying ChatGPT-4o, had around 1.76 trillion parameters^13^ and able to handle up to 111,616 tokens (approximately equates to 83,000–111,000 English words). When prompted on the number of tokens and parameters, Google Gemini 1.5 Pro had around 200 billion parameters and was able to handle up to 2,097,152 input tokens. Both ChatGPT-4o and Google Gemini 1.5 Pro were able to access metadata of PubMed abstracts nut unable to directly access the PubMed database.

### Scientific-focused large language model

For scientific-focused large language models (LLMs), three types of GPT-based systems were utilized:

1. LLMs with access to dedicated databases of full-length articles: These include models like GPT-4-based SciSpace (Typeset.io) and ScholarAI.
2. LLMs without dedicated databases but with direct access to publicly available sources: Examples include GPT-4-based ScholarGPT, which can access databases such as PubMed, arXiv, bioRxiv, Google Scholar, and patent databases.
3. LLMs without access to full-length articles, customized to draw information from abstracts: An example is PubMedGPT, which primarily utilizes abstracts from the PubMed database.

GPT-4-based SciSpace: SciSpace’s general knowledge cutoff was October 2023. However, it can retrieve, analyze, and synthesize information from its proprietary database of over 300 million research papers. Through its integration with this database, SciSpace provides real-time access to research papers up to 2025. Additionally, SciSpace can access publicly available databases, including CrossRef, PubMed, arXiv, Springer, Elsevier, IEEE Xplore, Google Scholar, and patent databases.

GPT-4-based ScholarAI: Like SciSpace, ScholarAI’s general knowledge cutoff was October 2023. It has access to the dedicated ScholarAI database, which contains more than 200 million research papers, and can also draw from publicly available sources such as CrossRef, PubMed, arXiv, Springer, Elsevier, IEEE Xplore, Google Scholar, and patent databases.

GPT-4-based ScholarGPT: ScholarGPT had a general knowledge cutoff in October 2023 and lacked a dedicated database. Instead, it relies on real-time access to abstracts and open-access full-text articles from publicly available sources, including CrossRef, PubMed, arXiv, Springer, Elsevier, IEEE Xplore, Google Scholar, and patent databases.

PubMedGPT: PubMedGPT was not trained on full-text manuscripts from PubMed or other databases. Its training included publicly available data up to 2023. PubMedGPT is designed to provide citations from PubMed and integrate findings from multiple studies to synthesize conclusions.

### Queries

Bone formation, fibrogenesis, cartilage formation, and cellular proliferation were utilized as proxies for phenotypes. These phenotypes were chosen as representative topics of interest for skeletal tissue researchers.

A power analysis was conducted to determine the minimum number of genes required to detect at least a 20% difference with a power of 80% and a significance level (p-value) of 0.05. The analysis indicated that at least 91 genes would be needed.

To ensure comprehensive representation, 102 genes previously reported in peer-reviewed studies to impact at least one of the four phenotypes were selected as genes of interest **(Table 1)**. These 102 genes were selected to encompass various genetic pathways such as bone morphogenetic pathway (BMP Pathway) (*BMP1, BMP2, BMP7, GREM1*, ***SMAD1***, *SMAD4, SMAD5, SMAD8, RUNX2, MSX2, PRX1*), wingless/integrated pathway (WNT Pathway) (*WNT1, WNT3A, LRP5, LRP6, RSPO1, RSPO2, RSPO3, DKK1, DKK2, DKK3, DKK4, SOST, FRZB*, CTNNB), the transforming-growth factor beta pathway (TGFβ pathway) (*TGFBR1, TGFBR2, CTGF, SMAD2, SMAD3, SMAD4, EGR1*), Hedgehog pathway (*GLI1, GLI2, GLI3*), NF-kB pathway (*NFKB1, IKBKB, RELA*), fibroblast growth factor pathway (FGF pathway) (FGF2, FGFR1, FGFR2, FGF23). For each gene, four distinct LLM queries were created, corresponding to the four phenotypes of interest: bone formation, fibrogenesis, cartilage formation, and cellular proliferation.

**Table 1.**
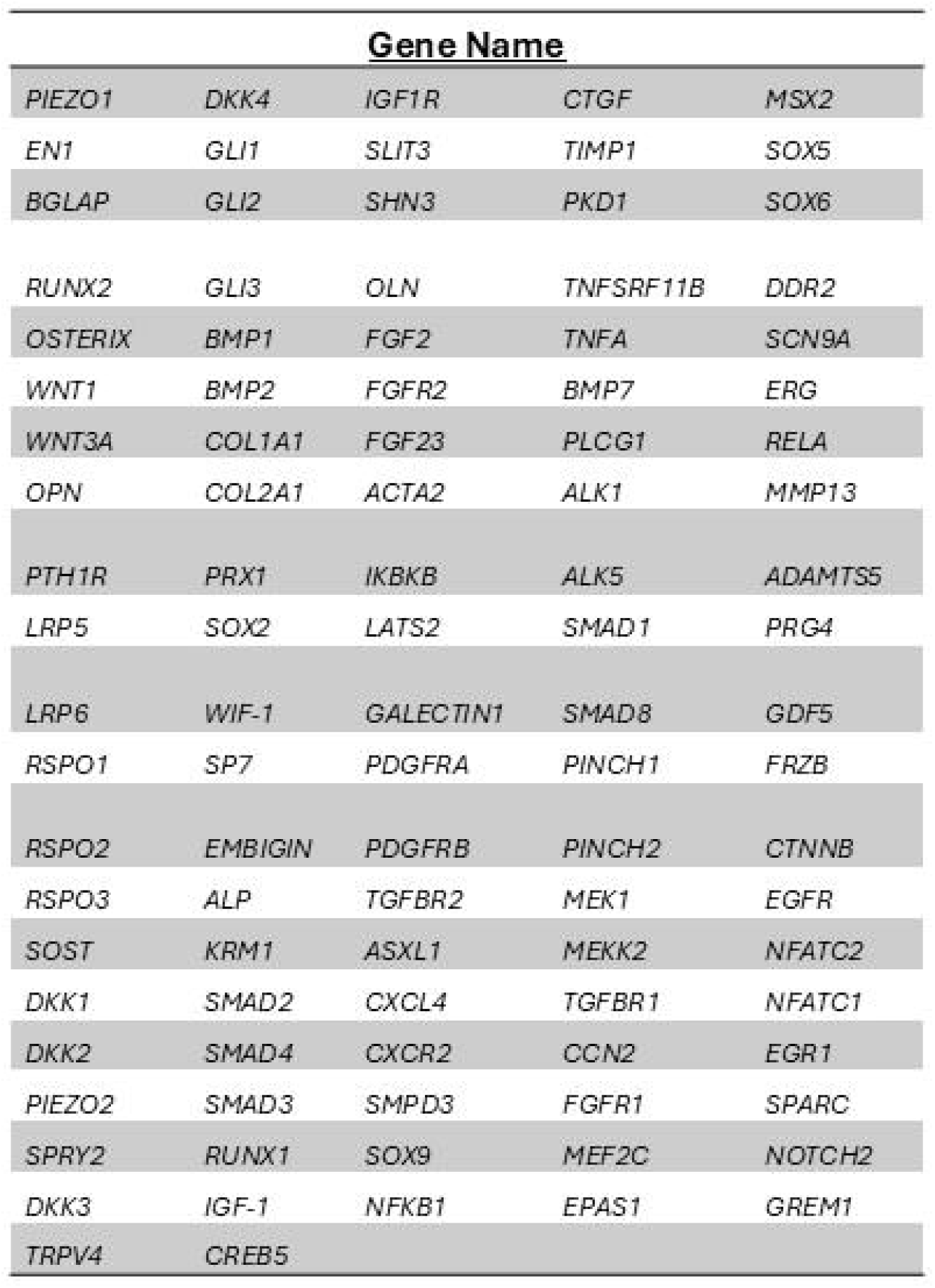
List of genes used in this study.

### LLM Prompts

The LLM prompt consisted of two sections. In the first section, the LLMs were prompted to decide, in one word, whether a gene showed an increase, decrease, or an inconclusive relationship with the queried phenotypes based on available peer-reviewed articles. In the second section, the LLMs were prompted to provide links to the peer-reviewed articles that informed their decisions.

The prompt was developed iteratively to ensure that the requested data were returned accurately and consistently. During initial testing, we observed that some LLMs occasionally provided links to articles that either did not exist or had titles unrelated to the query. Upon further prompting, the LLMs recognized these inconsistencies and revised their answers. We found that explicitly specifying the desired answer formats significantly improved the consistency of data extraction.

Does {genes[i]} increase or decrease {questions[j]} according to available peer reviewed papers?

In the first line of your answer, please answer with only 1 word: increase/decrease/inconclusive surrounded by a pair of square bracket.

In the following line of answers, please provide the links of paper that support your answer.

If you answered inconclusive, you don’t need to provide any links.

Here’s an example format: [increase / decrease / inconclusive]

Reference

1. https://pubmed.ncbi.nlm.nih.gov/123456/
2. https://www.mdpi.com/1234-4567/12/3/4567

The final prompt used in the study was as follows:

All links to peer-reviewed manuscripts provided by the LLM prompts were manually verified to ensure they corresponded to the intended and relevant manuscripts.

### Statistics

Logistic regression was performed to analyze the performance of large language models (LLMs) in providing correct answers, using human responses as the ground truth. ChatGPT-4o (base model) was treated as the reference model. The p-value was calculated to test the null hypothesis that the log-odds difference between the tested model and the reference model is zero (i.e., the performance of the tested model is the same as that of the reference model). Statistical analysis was conducted using GraphPad Prism v9.5, with a significance threshold set at p<0.05.

## Results

### ChatGPT-4o and Google Gemini 1.5 Pro demonstrated comparable moderate accuracy in mapping genes to phenotype

We investigated the performance of a OpenAI-based LLM (ChatGPT-4o) in comparison to the Google-based LLM (Google Gemini 1.5 Pro) to map genes to phenotype based on peer-reviewed literatures. To evaluate the ability of the LLMs to map genes to phenotypes, each model was tasked with mapping 102 genes (as determined by the power analysis described in the Methods section) to four phenotypes: bone formation, cartilage formation, fibrosis, and cell proliferation. This resulted in a total of four queries per gene and 412 queries per LLM. Manual mapping of genes to phenotypes by the senior author (VJS), based on peer-reviewed journal articles, was used as the ground truth for comparison.

ChatGPT-4o, demonstrated comparable accuracy in mapping of genes to phenotypes compared to the Google Gemini 1.5 Pro (**58.7%** for ChatGPT-4o and **58.2%** for Gemini 1.5 Pro, p=0.99, **Figure 1a**). Despite the one shot training explicitly querying for peer-reviewed articles, not all queries resulted in links to peer-reviewed articles. ChatGPT-4o provided significantly more links to non-peer reviewed articles than Gemini 1.5 Pro (**18.0%** for ChatGPT-4o and **6.7%** for Gemini 1.5 Pro, p=0.0001, **Figure 1b**). Additionally, ChatGPT-4o returned a higher percentage of irrelevant references compared to Gemini 1.5 Pro (**25.8%** for ChatGPT-4o and **17.5%** for Gemini 1.5 Pro, p=0.026, **Figure 1b**). Among the four queried phenotypes, ChatGPT-4o exhibited variable performance, achieving its highest accuracy in the fibrosis phenotype (72%) and its lowest accuracy in the cartilage formation phenotype (45%) (**Supplementary Figure 1**). Gemini 1.5 Pro, on the other hand, achieved similar performance across the four queried phenotypes, achieving its highest accuracy in the

**Figure 1.**
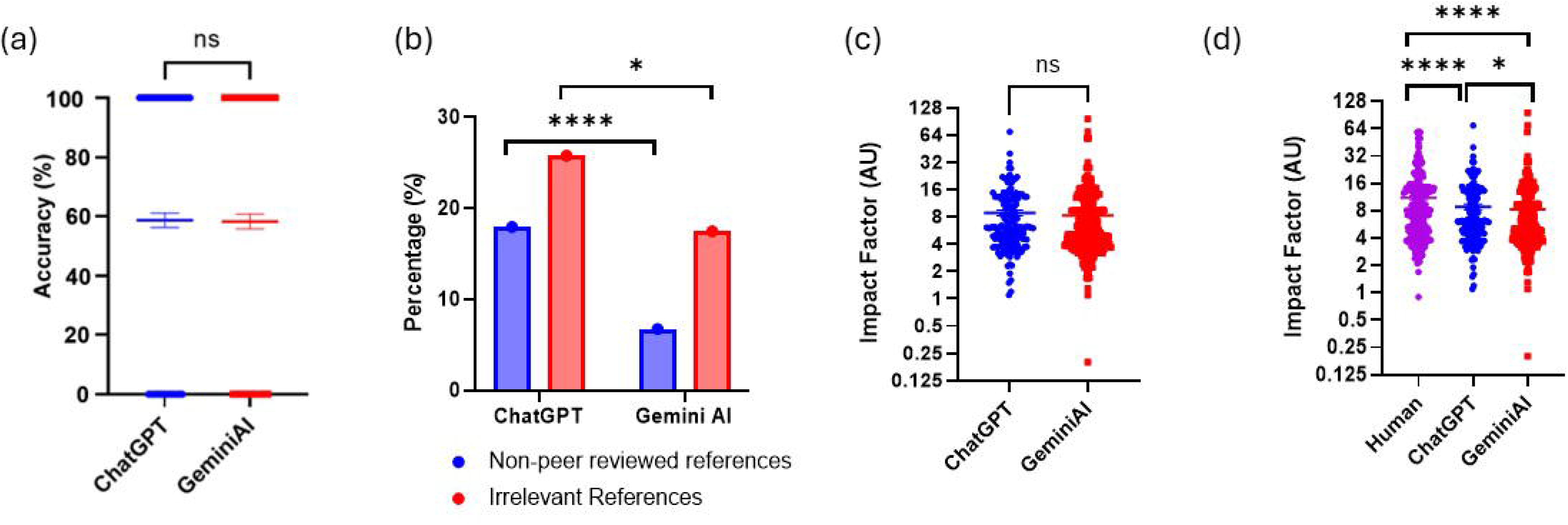
Accuracy and impact factor of references provided by ChatGPT4o and GeminiAI. **(a)** Accuracy of correct gene-to-phenotype mappings by ChatGPT4o and GeminiAI. Each dot represents a single gene-to-phenotype query, with a value of 1 indicating agreement between the tested LLM and human answers, and 0 indicating a discrepancy. Data are presented as mean± s.e.m., analyzed using the Wilcoxon pair-matched signed-rank test. **(b)** Proportion of non-peer-reviewed references (blue bars) and irrelevant references (red bars) provided by ChatGPT4o and GeminiAI. Data are presented as mean± s.e.m., analyzed using the Chi-squ are test.(c-d) Impact factor of references provided by human, ChatGPT4o, and GeminiAI. Each dot represents the impact factor of a single gene to-phenotype query. Data are presented as mean± s.e.m., analyzed using the Wilcoxon pair-matched signed-rank test. *p<0.05, **p<0.01, ***p<0.001, ****p<0.0001.

Among the relevant peer-reviewed articles and using the Wilcoxon pair-matched signed rank test, ChatGPT-4o cited references with a statistically significantly higher impact factor compared to Gemini 1.5 Pro (**9.0 ± 7.9** for ChatGPT-4o and **8.3 ± 9.6** for Gemini 1.5 Pro, **p = 0.042**; **Figure 1c**). Human-provided references had a statistically significantly higher impact factor than both ChatGPT-4o and Gemini 1.5 Pro (**p = 0.0001** for human vs. ChatGPT-4o, and **p = 0.0001** for human vs. Gemini 1.5 Pro; **Figure 1d**).

### The scientific-focused GPT-based LLM without its own database underperformed the base GPT-based LLM in mapping genes to phenotype

We investigated whether LLMs fine-tuned for scientific purposes could improve gene-phenotype mapping performance. To address this question, we utilized publicly available ChatGPT-4o-based, scientifically fine-tuned LLMs, including PubmedGPT, SciSpace, ScholarGPT, and ScholarAI. These scientific LLMs were categorized into three main groups: First, LLMs with access to dedicated databases of full-length articles (SciSpace and ScholarAI). Second, LLMs without dedicated databases but with direct access to publicly available sources (ScholarGPT). Third, LLMs without access to full-length articles, customized to extract information from PubMed abstracts (PubmedGPT).

PubmedGPT, an LLM customized to extract information from PubMed abstracts, demonstrated lower accuracy compared to ChatGPT-4o (**51.0%** for PubmedGPT vs **58.7%** for ChatGPT-4o, **p = 0.0014**; **Figure 2a**). However, PubmedGPT reported a significantly lower percentage of non-peer-reviewed references compared to ChatGPT-4o (**2.3%** vs. **18.0%**, respectively; **p = 0.0001**; **Figure 2b**). Despite this, PubmedGPT returned a higher percentage of irrelevant references compared to ChatGPT-4o (**63.6%** vs. **25.8%, p = 0.0001**; **Figure 2b**). Among the relevant peer-reviewed articles and using the Wilcoxon pair-matched signed rank test, the impact factor of references provided by PubmedGPT had significantly lower impact factors to those from ChatGPT-4o and human (**8.1 ± 9.7** for PubmedGPT, **8.9 ± 7.9** for ChatGPT-4o, **11.17 ± 10.5** for human, **p = 0.04** for PubmedGPT vs ChatGPT-4o and **p=0.0001** for PubmedGPT vs Human; **Figure 2c-d**).

**Figure 2.**
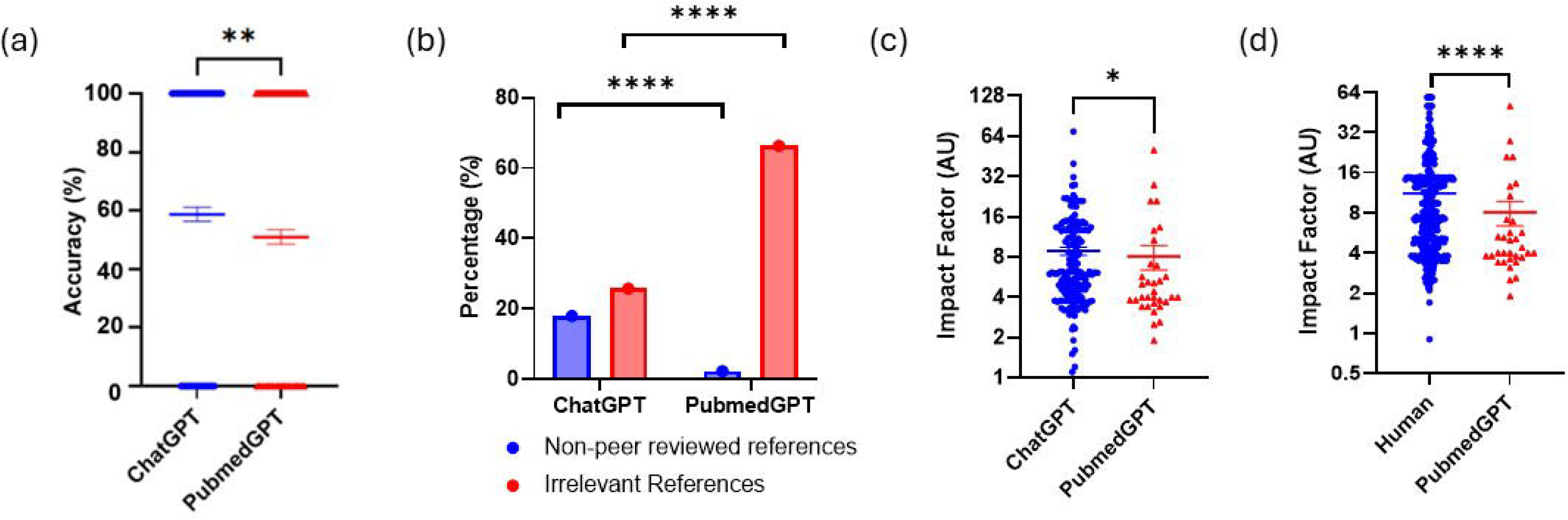
Accuracy and impact factor of references provided by PubmedGPT. **(a)** Accuracy of gene-to phenotype mappings by ChatGPT4o and PubmedGPT. Each dot represents a single gene-to-phenotype query, with a value of 1 indicating agreement between the tested LLM and human answers, and 0 indicating a discrepancy. Data are presented as mean± s.e.m., analyzed using the Wilcoxon pair-matched signed-rank test. **(b)** Proportion of non-peer-reviewed references (blue bars) and irrelevant references (red bars) provided by ChatGPT4o and PubmedGPT. Data are presented as mean± s.e.m., analyzed using the Chi-square test.(c-d) Impact factor of references provided by human, ChatGPT4o, and PubmedGPT. Each dot represents the impact factor of a single gene-to-phenotype query. Data are presented as mean± s.e.m., analyzed using the Wilcoxon pair-matched signed-rank test. *p<0.05, **p<0.01, ***p<0.001, ****p<0.0001.

### The scientific-focused GPT-based LLM without its own database underperformed the base GPT-based LLM in mapping genes to phenotype

To investigate whether GPT-based LLM with direct access to publicly available sources and databases (Pubmed, Arxiv, Biorxiv), we compared the ability of ScholarGPT to the base ChatGPT-4o and human to map genes to phenotype based on relevant peer-reviewed articles.

ScholarGPT had comparable accuracy compared to ChatGPT-4o (**52.9 %** for ScholarGPT vs **58.7%** for ChatGPT-4o, **p = 0.055**; **Figure 3a**). While all references provided by ScholarGPT are from peer-reviewed journals, 21.1 % of references provided are irrelevant to the particular gene and phenotypes queried, which was comparable to the base ChatGPT-4o (**Figure 3b**). Among the relevant peer-reviewed articles provided and using the Wilcoxon pair-matched signed rank test, the impact factor of references provided by ScholarGPT were comparable to the base ChatGPT-4o (**8.9 ± 7.9** for ChatGPT-4o, **8.75 ± 7.0** for ScholarGPT, **p = 0.70, Figure 3c**) but significantly lower compared to human (**8.75 ± 7.02** for ScholarGPT and **11.17 ± 10.5** for human, **p= 0.0001, Figure 3d**).

**Figure 3.**
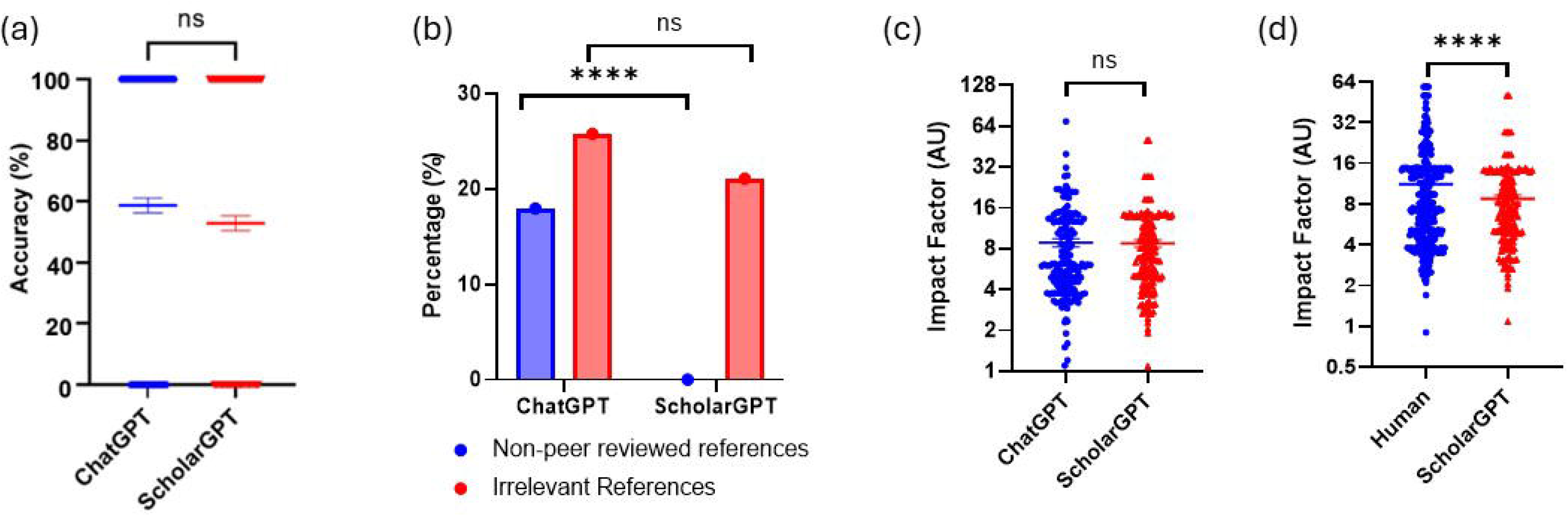
Accuracy and impact factor of references provided by ScholarGPT. **(a)** Accuracy of gene-to phenotype mappings by ChatGPT4o and ScholarGPT. Each dot represents a single gene-to-phenotype query, with a value of 1 indicating agreement between the tested LLM and human answers, and 0 indicating a discrepancy. Data are presented as mean± s.e.m., analyzed using the Wilcoxon pair-matched signed-rank test. **(b)** Proportion of non-peer-reviewed references (blue bars) and irrelevant references (red bars) provided by ChatGPT4o and ScholarGPT. Data are presented as mean± s.e.m., analyzed using the Chi-square test.(c-d) Impact factor of references provided by human, ChatGPT4o, and ScholarGPT. Each dot represents the impact factor of a single gene-to-phenotype query. Data are presented as mean± s.e.m., analyzed using the Wilcoxon pair-matched signed-rank test. *p<0.05, **p<0.01, ***p<0.001, ****p<0.0001.

**Figure 4.**
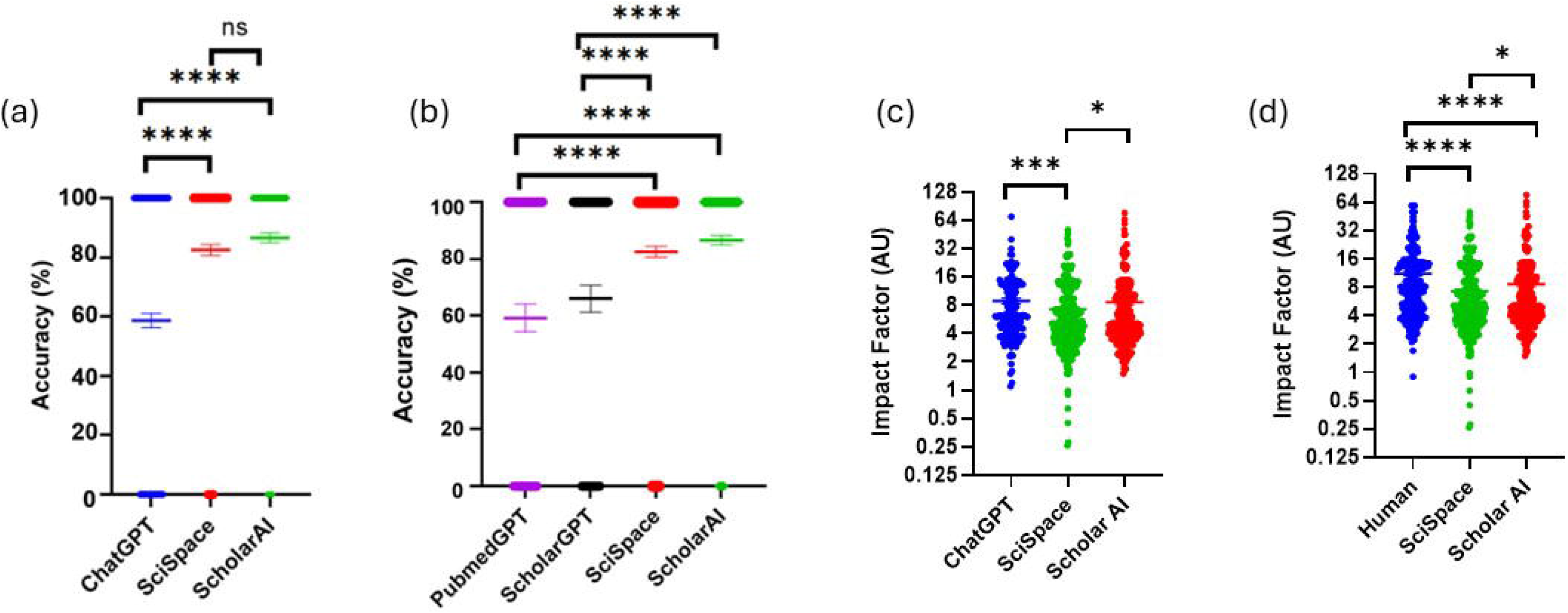
Accuracy and impact factor of references provided by SciSpace and ScholarAI. **(a)** Accuracy of gene-to phenotype mappings by ChatGPT4o, SciSpace, and ScholarAI. Each dot represents a single gene-to-phenotype query, with a value of 1 indicating agreement between the tested LLM and human answers, and 0 indicating a discrepancy. Data are presented as mean± s.e.m., analyzed using the Wilcoxon pair-matched signed-rank test. **(b)** Percentage of correct gene-to-phenotype mappings by PubmedGPT, ScholarGPT, SciSpace, and ScholarAI. Each dot represents a single gene to-phenotype query, with a value of 1 indicating agreement between the tested LLM and human answers, and 0 indicating a discrepancy. Data are presented as mean± s.e.m., analyzed using the Wilcoxon pair-matched signed-rank test. **(c-d)** Impact factor of references provided by human, ChatGPT 4o, SciSpace, and ScholarAl. Each dot represents the impact factor of a single gene-to-phenotype query. Data are presented as mean± s.e.m., analyzed using the Wilcoxon pair matched signed-rank test. *p<0.05, **p<0.01, ***p<0.001, ****p<0.0001.

### The scientific-focused GPT-based LLM with its own dedicated database outperformed the base GPT-based LLM in mapping genes to phenotype

We then investigated whether large language models (LLMs) with dedicated databases could enhance gene-phenotype mapping performance. LLMs equipped with dedicated databases of full-length articles, such as SciSpace and ScholarAI, demonstrated significantly higher accuracy in mapping genes to phenotypes compared to ChatGPT-4o (**82.5%** for SciSpace vs. **58.7%** for ChatGPT-4o, p = 0.0001; **86.7%** for ScholarAI vs. **58.7%** for ChatGPT-4o, **Figure 3a**). Moreover, SciSpace and ScholarAI outperformed LLMs without dedicated databases, such as those relying on publicly available sources (ScholarGPT), and LLMs tailored to retrieve information from PubMed abstracts (PubmedGPT) (**Figure 3b**).

Both SciSpace and ScholarAI exclusively provided references from peer-reviewed journals that were relevant to the queries (**Figure 3c**). Among the relevant references, ScholarAI produced references with impact factors comparable to those generated by ChatGPT-4o (**8.9 ± 7.9** for ChatGPT-4o vs. **8.6 ± 9.4** for ScholarAI; **p = 0.06, Figure 3c**). In contrast, SciSpace provided references with lower impact factors compared to ChatGPT-4o (**8.9 ± 7.9** for ChatGPT-4o vs. **7.2 ± 6.7** for SciSpace). The impact factors of references provided by ScholarAI were significantly higher than those provided by SciSpace (**8.6 ± 9.4** for ScholarAI vs. **7.2 ± 6.7** for SciSpace; **p = 0.047, Figure 3c**).

Overall, the impact factors of references supplied by SciSpace and ScholarAI were lower than those provided by human experts (**11.17 ± 10.5** for humans, **8.6 ± 9.4** for ScholarAI, and **7.2 ± 6.7** for SciSpace, **p=0.0001** for human vs ScholarAI and human vs SciSpace, **Figure 3d**).

## Discussion

Accurate gene-to-phenotype mapping at the cellular and tissue levels is critical for single-cell omics analysis^14^, understanding disease correlations^15^, assessing the effects of genetic perturbations^2^, and advancing the mechanistic understanding of disease pathophysiology^16^. Currently, human curation of gene-to-phenotype mappings based on peer-reviewed literature remains the gold standard. However, with the exponential growth in scientific publications, there is an urgent need for a minimally human-supervised system capable of performing this task with high accuracy.

The core requirements for such a system include: the ability to accurately retrieve all relevant literature, the capacity to extract and synthesize the desired information from the retrieved texts, and the capability to generate precise gene-to-phenotype correlations through aggregate analysis of the gathered data. Meeting these requirements would significantly enhance the efficiency and accuracy of gene-to-phenotype mapping while reducing the reliance on time-intensive manual curation. In this study, we evaluated the feasibility of leveraging transformer-based large language models (LLMs) to retrieve relevant literature and perform gene-to-phenotype mapping in an unsupervised manner. We then compared the accuracy and quality of retrieved literature to the gold standard human-curated gene-to-phenotype mappings. Additionally, we assessed the performance of fine-tuned LLMs, using different fine-tuning methods to determine which approach yielded the most accurate mappings.

Transformer-based large language models (LLMs) are increasingly recognized as promising and transformative tools in healthcare, clinical research, and basic science. These models demonstrate remarkable capabilities, such as passing the United States Medical Licensing Examination (USMLE)^17^ and excelling in medical subspecialty exams, often performing at or above the level of human experts^18-20^. However, despite the growing enthusiasm, these LLMs are not without risks and limitations, including the potential to misinterpret information or generate fabricated content. Bhattacharyya M et al. (2023) found that GPT-3.5 had very high rate of literature search inaccuracy (46%) and confabulation (47%)^21^. Walters WH et al. (2023) found that 55% of GPT-3.5 citation and 18% of GPT-4 citations are fabricated^22^. Chelli et al. (2024) demonstrated that LLMs such as GPT-3.5, GPT-4, and Bard exhibit high hallucination rates when tasked with extracting relevant peer-reviewed literature, with hallucination rates of 40% for GPT-3.5, 29% for GPT-4, and 91.4% for Bard^23^. These findings align with our results, which revealed that ChatGPT-4o returned 25.8% irrelevant literature and 18.0% non-peer-reviewed references.

ChatGPT-4o achieved an accuracy of only 58.7% in gene-to-phenotype mapping and generated references with significantly lower impact factors compared to manual literature searches. These findings align with the results of Chelli et al. (2024), who reported that even when using only appropriate references, GPT-3.5, GPT-4, and Bard failed to produce a comprehensive list of peer-reviewed references compared to standard database queries^23^.

To determine whether the poor accuracy and high hallucination rate observed in ChatGPT-4o is unique to this model, we tested another decoder-only transformer-based LLM, Gemini 1.5 Pro. Interestingly, while Gemini 1.5 Pro demonstrated comparable performance to ChatGPT-4o in mapping accuracy, it produced a significantly higher proportion of peer-reviewed references and a significantly lower rate of inaccurate references.

A likely explanation for the poor accuracy is the inability of GPT-4o and Gemini 1.5 Pro to directly access commonly used peer-reviewed article databases, such as PubMed and Unpaywall. This limitation restricts their ability to retrieve relevant peer-reviewed articles and synthesize the information into accurate gene-to-phenotype mappings^24^. Moreover, as LLMs, GPT-4o and Gemini 1.5 Pro are primarily designed for language processing rather than information processing, making them more prone to fabricating information (a phenomenon commonly referred to as “hallucination”)^22^.

To overcome the limitations of base GPT language models (LLMs) in conducting accurate and comprehensive literature searches, several specialized scientific LLMs have been developed. These models incorporate advanced features specifically designed to enhance the accuracy, depth, and reliability of literature retrieval and analysis. These enhancements include restricting the GPT to extract information solely from PubMed abstracts (e.g., PubMedGPT), providing the GPT with access to PubMed and other open-source databases (e.g., ScholarGPT), and augmenting the base GPT with proprietary databases containing hundreds of millions of full-length articles (e.g., ScholarAI and SciSpace).

Interestingly, the performance of specialized GPT models varied significantly depending on the type of specialization provided. For example, GPT-4-based PubMedGPT, which was fine-tuned to obtain information specifically from PubMed, demonstrated mixed results. PubMedGPT has the ability to access PubMed abstracts and metadata, and for articles labeled as open access, it can retrieve the full text through PubMed Central (PMC). While PubMedGPT ensured that all its references were sourced from peer-reviewed journals, 63.6% of the retrieved journals were irrelevant to both the gene name and the queried phenotypes—representing the highest irrelevance rate among all LLMs tested in this study. Furthermore, PubMedGPT underperformed all other LLMs, achieving only 51% accuracy in gene-to-phenotype mapping.

In contrast, GPT models with direct access to extensive repositories of over 200 million full-length articles, such as ScholarAI and SciSpace, demonstrated markedly superior performance in gene-to-phenotype mapping. These models consistently achieved an accuracy exceeding 80%, significantly outpacing other tested LLMs. Furthermore, all references provided by these models were not only peer-reviewed but also directly relevant to the queried gene-to-phenotype correlations, ensuring a higher level of reliability and precision in their outputs.

The enhanced performance of these models highlights the pivotal role of comprehensive database access in advancing the capabilities of large language models for specialized scientific tasks. By leveraging the wealth of information contained in full-length articles, rather than relying solely on abstracts or metadata, models like ScholarAI and SciSpace are better equipped to extract, synthesize, and contextualize complex relationships between genes and phenotypes. This level of access and integration enables them to overcome the limitations observed in other LLMs, such as irrelevant references or reduced mapping accuracy, making them invaluable tools for accelerating research in genomics and beyond. These findings align with prior studies demonstrating the superior performance of LLMs with database access in scientific research tasks compared to their non-fine-tuned counterparts.^13,25^

There are several limitations to this study. First, as of the time of writing this manuscript, there are over 20 scientific research-fine-tuned GPT models, each utilizing different fine-tuning methods and parameters. Second, our analysis sampled only 102 genes from a pool of more than 30,000 human genes^26^, alongside the vast number of potential phenotypes.

## Conclusion

This study represents the first systematic and comprehensive analysis of both base transformer-based large language models (LLMs) and fine-tuned LLMs built on the GPT-4 architecture. Specifically, we evaluated their ability to retrieve query-relevant scientific literature and synthesize this information to generate accurate gene-to-phenotype mappings. The GPT-4-based LLMs, particularly when equipped with direct access to databases, demonstrated remarkable performance in accurately mapping gene-to-phenotype. This integration not only enabled the generation of highly accurate gene-to-phenotype mappings but also ensured that the outputs were supported by relevant, peer-reviewed scientific articles. By combining cutting-edge natural language processing capabilities with access to high-quality data sources, GPT-4-based LLMs highlight their potential as powerful tools for facilitating complex biomedical research tasks, bridging the gap between vast genomic datasets and actionable biological insights.

## Author Contributions

Nicolas Suhardi and Vincentius Suhardi selected the list of genes and phenotypes, tested the relevance and accuracy of the references provided by the LLMs, conducted data analysis, and formulated questions. Anastasia Oktarina recorded the responses generated by all LLMs and verified the relevance and accuracy of the references provided by LLMs. Vincentius Suhardi performed the manual curation of references and manual gene-to-phenotype mapping that are used as ground truth for this study. Mathias Bostrom and Xu Yang assisted with editing the manuscript.

## Data Availability

The dataset used or analyzed in this study is available from the corresponding author upon reasonable request.

